# Model-Based Closed-Loop Control of Thalamic Deep Brain Stimulation

**DOI:** 10.1101/2023.12.20.572569

**Authors:** Yupeng Tian, Srikar Saradhi, Edward Bello, Matthew Johnson, Gabriele D’Eleuterio, Milos R. Popovic, Milad Lankarany

**Author notes:** To whom correspondence should be addressed: Milad Lankarany, The Krembil Research Institute – University Health Network (UHN), 60 Leonard Ave, Toronto, M5T 0S8, Canada.

## Abstract

Closed-loop control of deep brain stimulation (DBS) is crucial for effective and automatic treatments of various neurological disorders like Parkinson’s disease (PD) and essential tremor (ET). Manual (open-loop) DBS programming solely based on clinical observations relies on neurologists’ expertise and patients’ experience. The continuous stimulation in open-loop DBS may decrease battery life and cause side effects. On the contrary, a closed-loop DBS system utilizes a feedback biomarker/signal to track worsening (or improving) patient’s symptoms and offers several advantages compared to open-loop DBS. Existing closed-loop DBS control systems do not incorporate physiological mechanisms underlying the DBS or symptoms, for example how DBS modulates dynamics of synaptic plasticity. In this work, we proposed a computational framework for development of a model-based DBS controller where a biophysically-reasonable model can describe the relationship between DBS and neural activity, and a polynomial-based approximation can estimate the relationship between the neural and behavioral activity. A controller is utilized in our model in a quasi-real-time manner to find DBS patterns that significantly reduce the worsening of symptoms. These DBS patterns can be tested clinically by predicting the effect of DBS before delivering it to the patient. We applied this framework to the problem of finding optimal DBS frequencies for essential tremor given EMG recordings solely. Building on our recent network model of ventral intermediate nuclei (Vim), the main surgical target of the tremor, in response to DBS, we developed a biophysically-reasonable simulation in which physiological mechanisms underlying Vim-DBS are linked to symptomatic changes in EMG signals. By utilizing a PID controller, we showed that a closed-loop system can track EMG signals and adjusts the stimulation frequency of Vim-DBS so that the power of EMG in [2, 200] Hz reaches a desired target. We demonstrated that our model-based closed-loop control system of Vim-DBS finds an appropriate DBS frequency that aligns well with clinical studies. Our model-based closed-loop system is adaptable to different control targets, highlighting its potential usability for different diseases and personalized systems.

## Introduction

Deep brain stimulation (DBS) is a standard therapy for various movement disorders, including Parkinson’s disease (PD) [1], essential tremor (ET) [2], and dystonia [3]. The thalamic ventral intermediate nucleus (Vim) is the primary surgical target of DBS for ET treatments. The stimulation frequency of clinical Vim-DBS for treating ET is usually chosen to be ≥ 130 Hz [4][5][6]. Currently in clinics, DBS parameters – typically, frequency, amplitude and pulse width – are usually manually tuned with a trial-and-error process, based on immediate clinical observations by neurologists [7][8][9]. Such manual DBS programming may be biased towards the neurologists’ expertise and patients’ experience, while requiring multiple clinical visits to test a large number of possible parameters, which costs time and induces stress in both patients and clinicians [7][8][9]. Additionally, manually programmed DBS delivers continuous DBS (cDBS) to the patient, which can cause side effects and exacerbate stimulation habituation [10][11][12]. Continuous stimulation can also decrease battery life, thus increasing patients’ burden caused by battery replacement surgeries or battery recharging process [13][14]. Hence, there is a need for a control system that can automatically adjust DBS parameters in a closed-loop fashion. Such closed-loop DBS needs to be based on a biomarker which characterizes the patient’s clinical states.

A closed-loop DBS control system consists of three essential components: (*i*) the input DBS pulses; (*ii*) the output feedback, i.e., the biomarker observed during DBS; and (*iii*) the feedback control, which adjusts the DBS parameters based on the feedback biomarker [15][16][9]. The closed-loop DBS systems offer automatic ways to adapt stimulation parameters moment-to-moment with respect to the patient’s clinical states [15]. Compared with the manually programmed (open-loop) cDBS, the closed-loop DBS can significantly reduce the stimulation time and enhances clinical efficacy (less side effects) [15][17][10].

Most common closed-loop DBS systems use local field potential (LFP), recorded from stimulated nuclei, to find an effective feedback biomarker [17][18][19], for example the power of the beta oscillation (12-32 Hz) of LFP recorded in the subthalamic nucleus (STN) for reducing PD symptoms [9][15][17]. Velisar et al. (2019) [19] developed a closed-loop DBS control system in which the beta power of the STN-LFP was chosen as the biomarker, and the DBS amplitude was updated by a dual-threshold control method that maintains the STN-LFP beta power to be within a certain range. Other signals like muscle activities in electromyography (EMG) or inertial measurement unit (IMU) have been also used as the biomarker in closed-loop DBS treatment of tremors [10][20][21][22]. For example, in the treatment of ET, Herron et al. (2017) [21] developed a closed-loop DBS system that controls the EMG power to be below a specified threshold. There are also other types of feedback biomarkers used in closed-loop DBS, e.g., single-unit recordings and the coherence among electroencephalogram (EEG) recordings [24].

Regardless of the type of the feedback biomarker, DBS setting is determined based on neural (e.g., LFP) or behavioral (e.g., EMG) signals in the most existing closed-loop DBS controllers [19][17][21][25][26][27]. However, these methods suffer from the lack of mechanistic understanding of the physiological mechanisms underlying the DBS and disease-related neuronal circuits. An effective approach to overcome this problem is to embed a computational model of the underlying mechanisms into the control system [9]. For example, to control Parkinson’s disease, closed-loop DBS systems were developed based on physiological models of the related cortico-basal ganglia-thalamic network, e.g., in Liu et al. (2021) [28] and Fleming et al. (2020) [29]. Liu et al. (2021) [28] used the control system to suppress the beta oscillations in cortex, and Fleming et al. (2020) [29] tried to suppress the beta power of LFP in STN. Although these computational studies included computational models for adjusting DBS in a closed-loop manner [29][9][28], the models were not validated for replicating/tracking experimental data nor they incorporated DBS mechanisms of actions, e.g., DBS-induced short-term synaptic plasticity [30][31][32].

In this work, we develop a closed-loop control system to adjust the stimulation frequency of Vim-DBS automatically. Our control system is based on a computational model that predicts the EMG activities in response to different frequencies of Vim-DBS. In this computational model, the firing rate of the Vim neurons in response to Vim-DBS is predicted by our previous rate network model that reproduced the human clinical data recorded in Vim neurons receiving DBS of different stimulation frequencies (10 – 200 Hz) [33]. Our rate network model incorporated the DBS effect on dynamics of short-term synaptic plasticity [33]. We use a biophysically-reasonable simulation study including models of DBS, Vim, motor cortex, motor neurons in the spinal cord, and muscle fibers to generate muscle activities (represented by EMG). To link Vim-DBS to EMG signals in our model-based control framework, model-predicted EMG signals, generated by our simulation study, are used to calculate the feedback biomarker by a polynomial fit, which is processed and implemented in a proportional-integral-derivative (PID) controller [34][35][36] that automatically updates the appropriate DBS frequency. Our model-predicted EMG can replicate the essential tremor symptoms during DBS-OFF, and is consistent with clinical observations of tremor during different frequencies of Vim-DBS. In a closed-loop DBS control system, the ability of predicting the biomarker decreases the probability of delivering inappropriate DBS frequencies to the patient, and thus increases therapeutic efficacy and reduces side effects.

We anticipate that our computational framework can facilitate the development of model-based control systems which can be potentially implemented in and out of the clinic to automatically update the appropriate DBS frequency for individual patients suffering from different diseases.

## Materials and Methods

We developed a computational framework for incorporating physiological mechanisms of deep brain stimulation into controlling disease symptoms. This framework consists of two main parts: (1) A computational model characterizing the physiological mechanism of the stimulated neuronal network and (structurally/functionally) connected neurons; and (2) A feedback control reflecting the disease state. In this study, we use computer simulations and apply our framework to control DBS frequency to reduce ET symptoms observed from EMG signals.

### Computational Model

The computational model consists of four components: (*i*) neural activities, spikes, generated by the Vim network model in response to different DBS frequencies; (*ii*) motor cortex neural activities influenced by propagated Vim-DBS effects to motor cortex; (*iii*) spinal motor neurons activities impacted by neurons in the motor cortex; (*iv*) motor unit action potentials in the muscle fibers innervated by the spinal motor neurons.

#### (i) Vim-network model impacted by Vim-DBS

The firing rate dynamics of the Vim neurons in response to Vim-DBS were simulated by our previous model of the Vim-network based on clinical DBS data recorded during surgery on human patients with essential tremor (ET) [33]. Our previous model could accurately reproduce the clinically recorded instantaneous firing rate of the Vim neurons receiving DBS of different stimulation frequencies (10 – 200 Hz) [33].

#### (ii) Propagation to the primary motor cortex

In our model, the Vim-DBS effects are propagated to the primary motor cortex (M1). We modeled the propagation to an M1 neuron of an ET patient using the dynamics induced by two sources: (a) the effects of Vim-DBS; and (b) the background neuronal activities that induce the tremor symptom. The Vim-DBS effects are from the direct DBS activation of the axons projected to the M1 neuron and the firings of the Vim neurons during Vim-DBS.

These effects consist of the direct axon activation and DBS-induced Vim firings. DBS activates the axons connecting to the synapses projecting to the M1 neuron, and these synapses are characterized by the Tsodyks & Markram (TM) model [37] (**Supplementary Method**). Besides the direct axon activation, the M1 neuron is also affected by the DBS-induced firings of the Vim neurons. With our previous Vim-network model [33], we simulated the instantaneous firing rate of the Vim neurons receiving DBS of different stimulation frequencies (10 – 200 Hz). The Vim firing rate signal is the time-varying Poisson rate for generating Poisson spike trains, which were passed to the TM-modeled synapses to produce the post-synaptic current in the M1 neuron (**Supplementary Method**).

In addition to the DBS effects, we also modeled the background neuronal activities inducing the tremor symptom. The tremor activities observed in the EMG from ET patients are often in the frequency band 4 – 8 Hz [38][39][21]. The tremor-inducing background firing rate was taken as a waveform consisting of 6-Hz bursts with a baseline shift (**Supplementary Figure 2**). To be consistent with the EMG recordings from ET patients [38][40][39][41], each burst consists of 3 consecutive sinusoidal waves and the period of each wave is 20ms (**Supplementary Figure 2**). We then generated Poisson spike trains from the background firing rate waveform; these spikes were then passed to the M1 synapses characterized by the TM model [37], which produced the post-synaptic current into the M1 neuron (**Supplementary Method**).

The membrane potential of the M1 neuron was characterized by a leaky integrate-and-fire (LIF) model as follows:

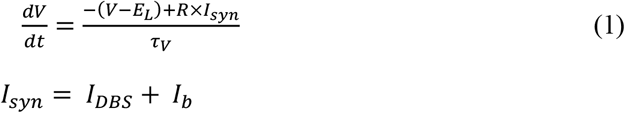

where *E*_*L*_= −65 mV is the equilibrium potential, R (resistance parameter) = 1 MΩ, and *τ*_*V*_ = 10 ms is the membrane time constant; spikes occur when *V* ≥ *V*_*th*_, where *V*_*th*_ = −35 mV. The reset voltage is −90 mV and the absolute refractory period is 1ms. *I*_*syn*_ is the total post-synaptic input current, consisting of the inputs induced by Vim-DBS (*I*_*DBS*_) and the background inputs generating the tremor (*I*_*b*_), and was obtained by the Tsodyks & Markram model [37] that incorporates all the input spikes (see **Supplementary Method**).

#### (iii) Projection from primary motor cortex to spinal motor neurons

We modeled the Vim-DBS effects as being propagated to a population of 150 M1 neurons (*N*_*c*_ = 150), which project to 120 motor neurons (*N*_*m*_ = 120) in the spinal cord [42][43]. We assumed that each motor neuron randomly connects to 70 M1 neurons, and receives monosynaptic inputs from each M1 neuron [42][44]. Following a spike from an M1 neuron, we modeled the post-synaptic current into a motor neuron by the rule of spike time dependent plasticity (STDP) from Izhikevich (2006) [45]:

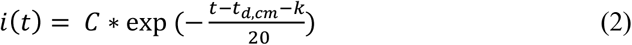

where *k* is a spike timing of an M1 neuron, *t*_*d,cm*_ = 10 ms is the M1-to-spinal-motor-neuron transmission delay [46], *t* ≥ *k* + *t*_*d,cm*_ is a time point following the M1 spike timing *k*, and *C* = 0.1 is the scaling factor [45]. The membrane potential of the motor neuron is given by an LIF model equivalent to that of Herrmann and Gerstner (2002) [47][42]:

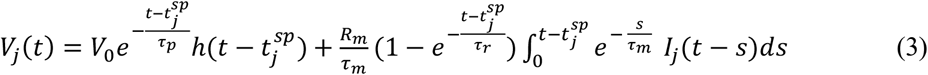

where 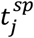 is the last spike timing of the j^th^ motor neuron. h() is the Heaviside step function, which is zero when its argument is negative and one otherwise. *I*_*j*_(*t*) is the post-synaptic current from M1 neurons into the j^th^ motor neuron, and is a summation of the STDP (**Equation 2**) from each of the 70 M1 neurons projecting to the motor neuron. *V*_0_ = −22 mV is the reset membrane potential [42]. When the membrane potential reaches a firing threshold *V*_*th*_, it is instantaneously reset to *V*_0_, and the integration process restarts. For each motor neuron, the firing threshold *V*_*th*_ ∈ [5, 15] mV is chosen randomly [42]. *R*_*m*_= 36 MΩ is the input resistance [42]. *τ*_*p*_ = 2 ms is the refractory time constant, *τ*_*m*_ = 4 ms is the passive membrane time constant, *τ*_*r*_ = 100 ms is the recovery time constant [48][42]. The firing rate of a human motor neuron is normally between 5 and 50 Hz [49], although it can be over 100 Hz for a brief period during fast contractions [50].

#### (iv) Generation of EMG activities from spinal motor neuron spikes

The spikes from the spinal motor neurons generate action potentials in the motor units of muscle fibers [42][43]. These motor unit action potentials (MUAP) were modeled as follows [43][42][51]:

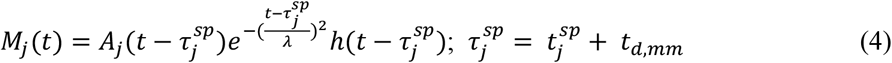

where *M*_*j*_ is the MUAP of the j^th^ motor unit, corresponding to the j^th^ motor neuron, *λ* = 2 ms is the time factor [42], 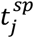 is the spike timing, *t*_*d,mm*_ = 10 ms is the motor-neuron-to-muscle conduction delay in human [52], and h() is the Heaviside step function. *A*_*j*_ is the scale factor of the amplitude of activities of the j^th^ motor unit [53][42], and was modeled as following the exponential distribution 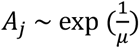, where *μ* is the mean of distribution [53][42]; we chose *μ* = 7 × 10^−3^ to be consistent with the EMG simulation during transcranial magnetic stimulation (TMS) in Moezzi et al. (2018) [42]. Finally, the surface EMG was modeled as the summation of MUAPs with a low-level Gaussian white noise [43][42].

### Feedback Control for DBS Frequency

Our computational model simulates the EMG signal in response to different DBS frequencies. Simulated EMG signals are used to calculate the feedback biomarker which controls the DBS frequency. The feedback control consists of three main parts: (1) biomarker identification, (2) computation of a system output from the biomarker, and (3) a closed-loop controller that implements the system output to update the DBS frequency.

#### Biomarker identification

The EMG simulation with our computational model is slow to implement: it takes more than 30 minutes using MATLAB R2022b with a personal computer to simulate 10 s of EMG signal. Thus, to facilitate the implementation speed of the model in a closed-loop control system, we need a fast EMG estimation to replace the direct EMG simulation. We implemented a polynomial method to estimate the EMG from the Vim firing rate, and the direct model simulated EMG is used as the reference (i.e., reference EMG) for estimation. We implemented the MATLAB custom function *polyfit* for the polynomial estimation, which gives a least-square fit of the polynomial coefficients. The estimated EMG is formulated as follows:

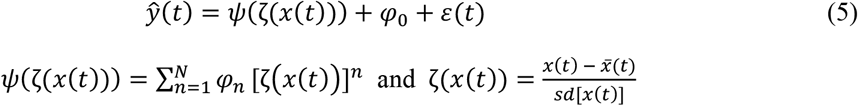

where *x*(*t*) is the Vim firing rate simulated from our previous Vim-network model [33], and *ŷ*(*t*) is the estimated EMG. *ζ*(*x*(*t*)) is the standardization of 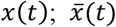 and *sd*[*x*(*t*)] are the mean and standard deviation of *x*(*t*), respectively. The polynomial order is 25 and {*φ*_0_, *φ*_1_, …, *φ*_*N*_, *N* = 25} are the polynomial coefficients. The *R*^2^ statistic [54] of the fit generally increases as the polynomial order increases (**Supplementary Figure 4**), and 25 is a minimal order when *R*^2^ converges. *ε*(*t*) is the Gaussian white noise with standard deviation = 0.038. We fitted the consistent polynomial coefficients (**Supplementary Table 2**) across data from different DBS frequencies, including {10, 50, 80, 100, 120, 130, 140, 160, and 200 Hz}. Our previous work showed that the consistent model parameters fitted based on concatenated DBS frequencies in a certain range – in this case, [10, 200] Hz – can be consistently applied to unobserved frequencies (e.g., 25 Hz and 180 Hz) in the same range [31]. Thus, we implement the polynomial estimated EMG as the biomarker to control the DBS frequencies in the range [10, 200] Hz.

#### Computation of system output from the estimated EMG

The power spectral density (PSD) of the estimated EMG *ŷ*(*t*) (**Equation 5**) is the system output for updating the frequency of the input DBS (10 – 200 Hz). In the system output, we consider PSD in the frequency band [2, 200] Hz, which includes the frequencies of both DBS and EMG activities [38][39][21][30]. The power density is calculated by the following equation [55][28], with sampling frequency 0.5 kHz:

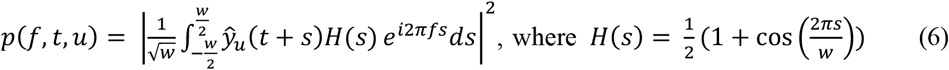

where *f* represents the frequency of the EMG activities. *H*(*s*) is the Hann windowing function [55][56], and *w* is the width of the window chosen to be w = 1 s to ensure stability [28][55]. *ŷ*_*u*_(*t*) represents the estimated EMG in response to DBS with stimulation frequency *u*. The total power of EMG activities over the initial period *T* of the estimated EMG signal is then given as:

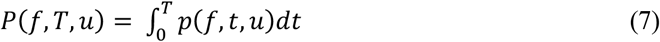

The EMG power thus computed can be used to analyze the EMG activities in the frequency domain in response to different frequencies of DBS.

In response to DBS frequency *u*, the PSD within the frequency band [2, 200] Hz of EMG activities is:

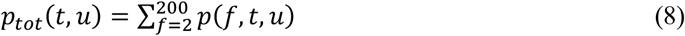

The system output *z*(*u*) is calculated as the mean PSD in the initial T = 5 s of the estimated EMG:

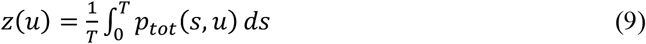

Finally, a closed-loop controller updates the DBS frequency *u* so that *z*(*u*) is close to a specified target value *β*_*z*_ (see the next section).

#### The PID controller that updates the DBS frequency

We implemented the proportional-integral-derivative (PID) controller [34][35][36] to update the DBS frequency based on the system output, *z* (**Equation 9**). The updated DBS frequency *u*(*t*) is computed as follows,

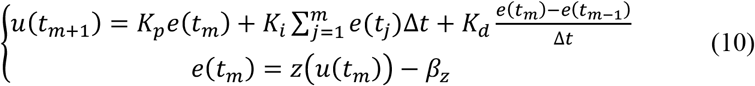

and is evaluated at the time points {*t*_1_, *t*_2_, …, *t*_*M*_}, with *t*_1_ = 0, *t*_*M*_ = 10 min, and ∆*t* = *t*_*m*_ - *t*_*m*−1_ = 1/3 min. The simulation of the PID controller was performed with MATLAB R2022b. *K*_*p*_, *K*_*i*_, and *K*_*d*_ denote the proportional, integral, and derivative gains, respectively. The error signal *e*(*t*) is the difference between the system output *z* and its target value *β*_*z*_.

## Results

Our proposed computational framework for model-based closed-loop DBS control is shown in **Figure 1**. In **Figure 1** (**A**), we utilize a biophysically-realistic encoding model to replicate (predict) neuronal activities in response to DBS (arbitrary) patterns. In this encoding model, the dynamics of synaptic plasticity and other biophysical details can be preserved and model parameters are estimated by fitting the model output to experimental data [33][31]. Additionally, to map neural activities to behavioral signals, we use data-driven approaches in our framework. A controller, e.g., a PID controller, is developed to adjust DBS patterns given behavioral signals solely. It is to be noted that, unlike encoding model, using biophysically-realistic models to identify neural-behavioral relationship requires building a network of several neuronal circuits (see our simulation study in **Figure 2**), which in turn increases the complexity of this type of models for mapping neural features to behavioral activities. However, we show that data-driven approaches are strong alternatives. We summarized the proposed computational framework in **Figure 1** (**B**) to highlight contributions of encoding (biophysical) and decoding (data-driven) models.

**Figure 1.**
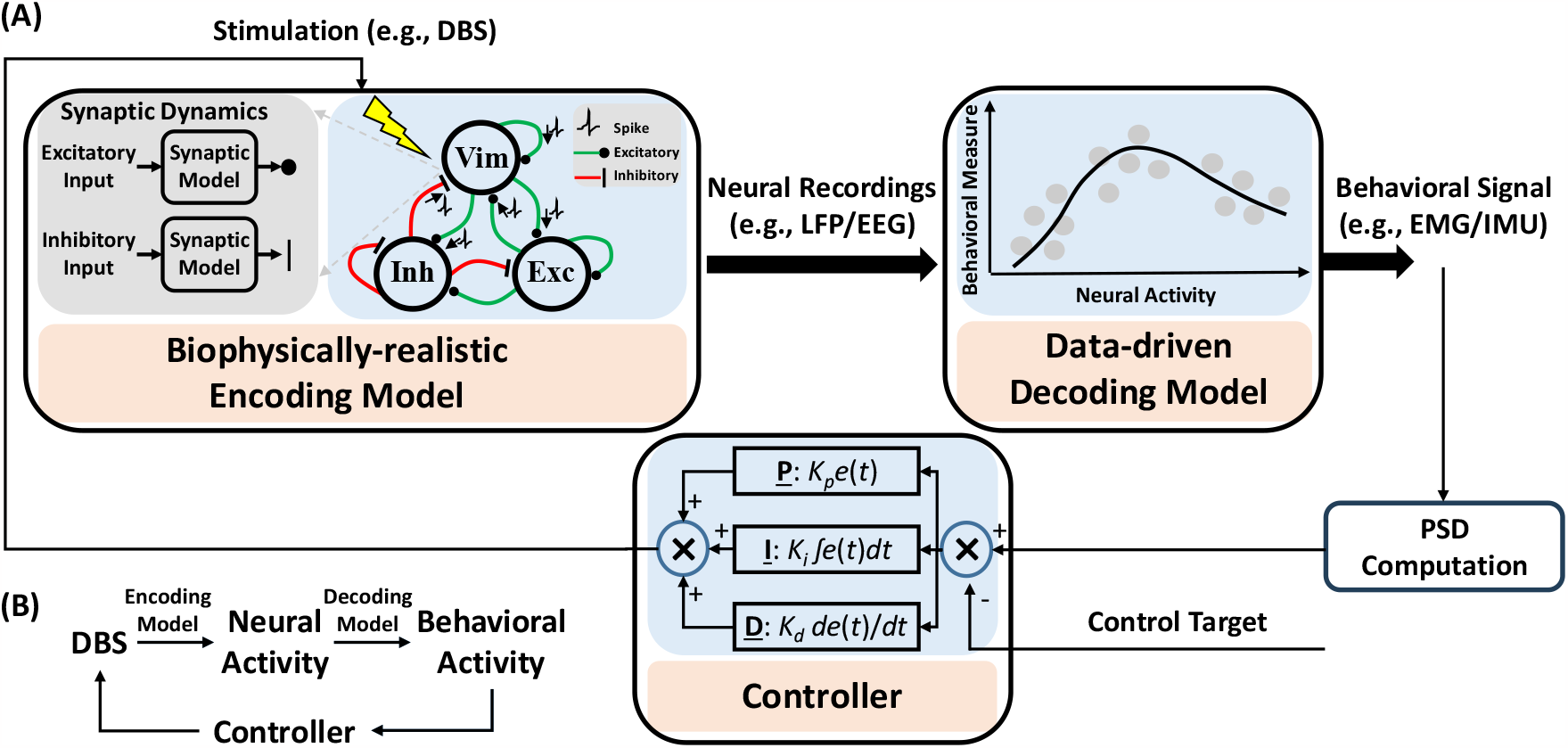
Computational Framework for Proposed Model-based Closed-Loop DBS Control System. (A) Biophysical details, e.g., dynamics of synaptic plasticity, are preserved in the encoding model to identify how different patterns of DBS change neural activities of stimulated neurons. A data-driven decoding model was used to map neural activity to behavioral signals like EMG. A controller is utilized to adjust DBS in a closed-loop manner. (B) A summary diagram of proposed computational framework.

**Figure 2.**
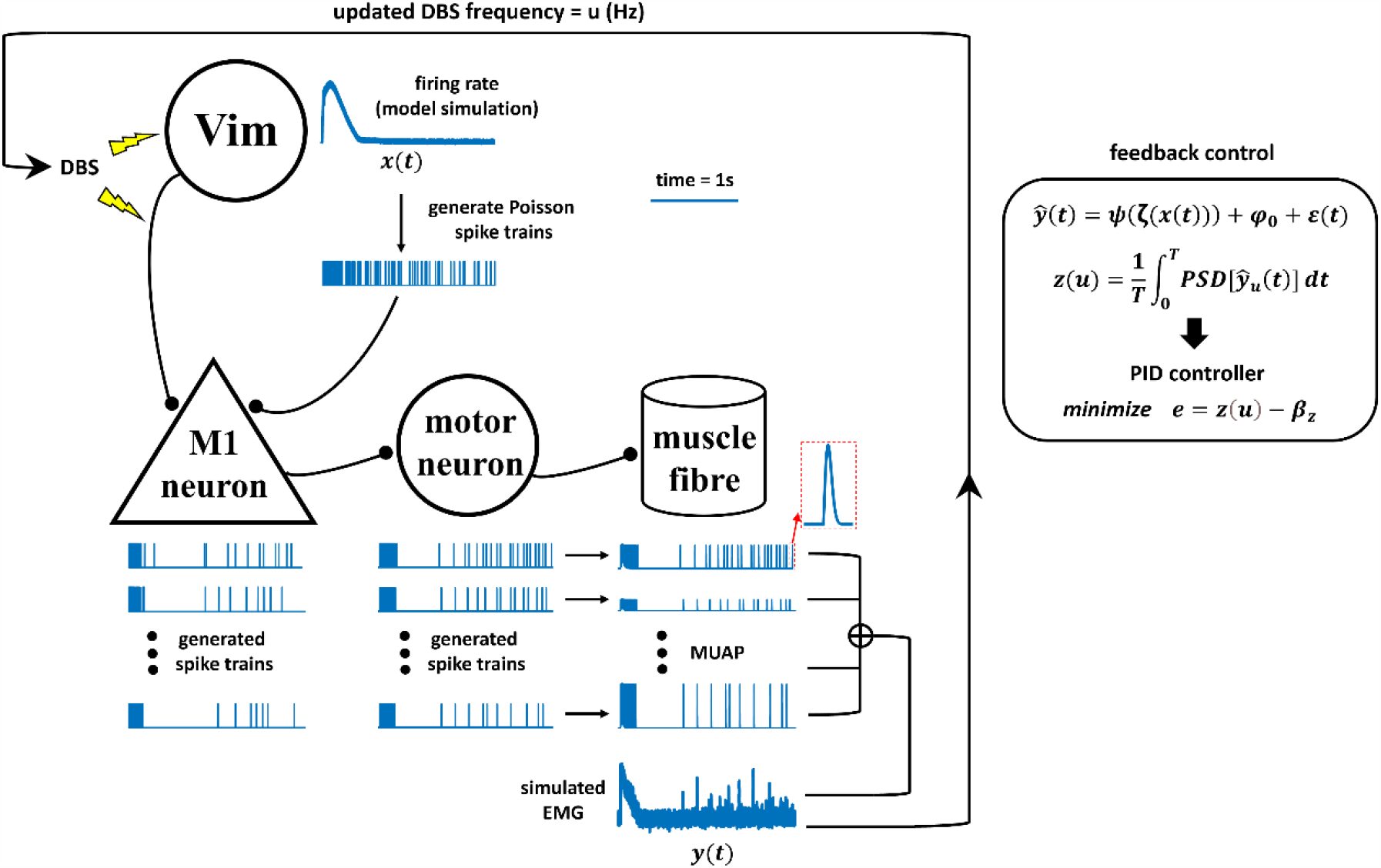
Schematics of biophysically-reasonable simulation study for Vim-DBS Control. DBS pulses are delivered to the neurons in the thalamic ventral intermediate nucleus (Vim), and also activate the axons projecting to neurons in the primary motor cortex (M1). The firing rate of the Vim neurons impacted by DBS was obtained from our previous Vim-network model that reproduced experimental DBS data [33]. The modeled Vim firing rate (x(t)) is also the Poisson rates for generating spike trains, which are propagated to the M1 neurons. The spikes from the M1 neurons are then propagated to the motor neurons in the spinal cord. The spikes from these motor neurons innervate the corresponding motor units in the muscle fibers, and induce motor unit action potentials (MUAP). The simulated EMG (y(t)) is a linear summation of MUAPs with additive low-level Gaussian white noise. In the feedback control, we estimate the simulated EMG by fitting a polynomial function ψ (**Equation 5**), and this estimated EMG (ŷ(t)) is the biomarker used in control. φ_0_ is a constant term of ψ, ε(t) is Gaussian white noise and ζ(x(t)) is the standardization of the modeled Vim firing rate x(t) (**Equation 5**). The system output z(u) is calculated as the mean power spectral density (PSD) over the initial T of the biomarker ŷ (t) (**Equation 9**). Finally, a proportional-integral-derivative (PID) controller (**Equation 10**) updates the DBS frequency u that reduces the error (e) between the system output z(u) and a specified target value β_z_.

In next sections, we present details of the model-based closed-loop DBS system which can effectively control the DBS frequency based on EMG signals generated by a biophysically-reasonable simulation (**Figure 2**) of the underlying neuronal network (from DBS/Vim neural activity to muscle activity).

### Schematics of biophysically-reasonable simulation study for Vim-DBS Control

The simulation study for Vim-DBS control system is schematized in **Figure 2**. The effects of Vim-DBS consisted of the direct axon activation and the DBS-induced Vim firings, which were simulated from our previous Vim-network model established based on clinical Vim-DBS data [33]. The Vim-DBS effects are propagated to the M1 neuron, which projects to the motor neuron in the spinal cord. The firings of the spinal motor neurons innervate the corresponding motor units in the muscle fibers, and induce motor unit action potentials (MUAP). The simulated EMG consists of a linear summation of MUAPs and a low-level Gaussian white noise. In the feedback control, in order to facilitate the implementation speed, we estimated the simulated EMG by a polynomial fit. Then we computed the mean power spectral density (PSD) of such polynomial-estimated EMG as the system output. Finally, a proportional-integral-derivative (PID) controller updated the DBS frequency that brought the system output close to a specified target value.

### The Vim-Cortex Propagation

Our model of the propagation of the Vim-DBS effects to the cortical M1 neurons was validated by using a 10-s block of neural spike data from non-human primate single-unit recordings in M1 during 130-Hz VPLo-DBS, reported in Agnesi et al. (2015) [57]. Thalamic VPLo nucleus (ventralis posterior lateralis pars oralis, in the Olszewski atlas) in non-human primate is homologous to the Vim human [58][59]. For quantifying the effect of 130-Hz DBS on spike activity, a peristimulus time histogram was derived based on the timing of spike events occurring between 0 to 7.7 ms after each DBS pulse event, and visualized by convolving the spikes with a Gaussian kernel (**Figure 3**), both for empirical and simulated data. The standard deviation of the Gaussian kernel was 0.2 ms, which was obtained from a method that optimized the Gaussian kernel to best characterize the spikes using a Poisson process [60][61][33]. Our Vim-M1 propagation model’s generated M1 spike activity behaved similarly to that of the empirically-recorded non-human primate M1, as indicated by spike-raster and PSTH firing rate analysis (**Figure 3**). Thus, our Vim-M1 propagation model could reflect the M1 dynamics during 130-Hz Vim-DBS.

**Figure 3.**
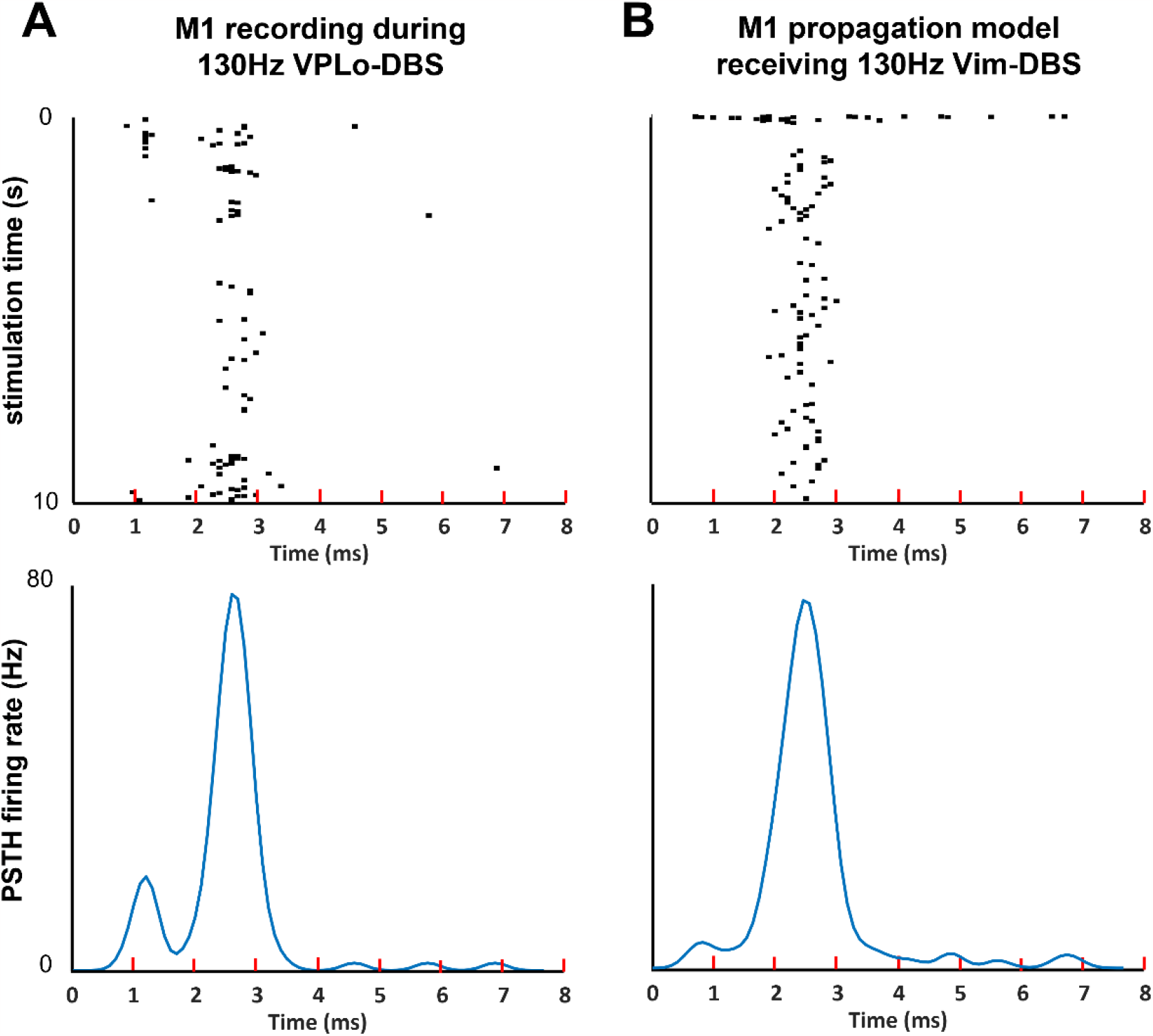
Raster Plot and PSTH of Model Simulation and Non-human primate Recording. (A – B) The spikes times occurring within each inter-pulse-interval during 10 seconds of 130-Hz DBS are visualized as a raster plot. We obtain an estimate of the instantaneous firing rate induced around these each DBS event by computing a peristimulus time histograms (PSTH), convolving the spikes with a 0.2 ms Gaussian kernel. (A)The raster plot and PSTH of the non-human primate single-unit recording in the primary motor cortex (M1) during 130-Hz VPLo-DBS [57]. (B)The raster plot and PSTH of our model simulation of M1 spikes during 130-Hz Vim-DBS.

### EMG Simulation from the Computational Model

We simulated EMG from our model, both during DBS-OFF and in response to Vim-DBS of different stimulation frequencies in [10, 200] Hz (**Figure 4**).

**Figure 4.**
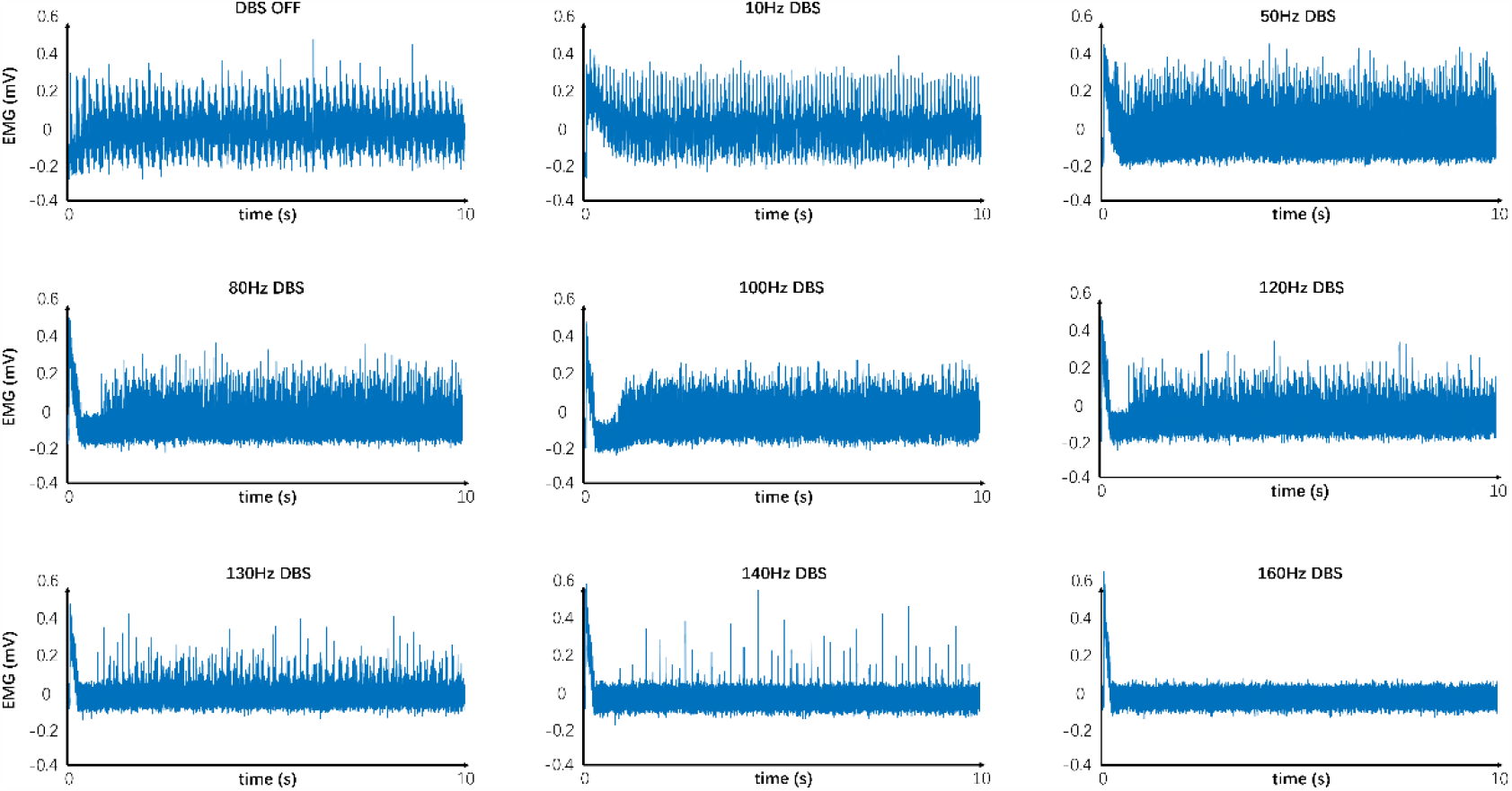
Model Simulated EMG in Response to Different Frequencies of DBS. Simulated EMG with our model, in response to different frequencies of Vim-DBS. Each signal is given relative to its mean.

The EMG simulation with DBS-OFF presented the typical tremor band (∼ 6 Hz) in the clinical EMG signals recorded from ET patients [38][39][21] (**Figure 4**). During low frequency (≤ 50 Hz) of DBS, the amplitude of the simulated EMG is similar to (or slightly higher than) the DBS-OFF situation (**Figure 4**). Such simulation is consistent with the clinical observations that low-frequency Vim-DBS (≤ 50 Hz) is often ineffective and can exacerbate the tremor [62][63][64]. The simulated EMG amplitude is lower than the DBS-OFF situation when DBS frequency is ≥ 80 Hz (**Figure 4**). During high frequency (≥ 100 Hz) of DBS, the simulated EMG amplitude is clearly depressed compared with the DBS-OFF situation (**Figure 4**). Such simulation is consistent with the clinical observations that high frequency (≥ 100 Hz) Vim-DBS can decrease the tremor [64][63][40]. The simulated EMG is mostly suppressed when DBS frequency is ≥ 130 Hz (**Figure 4**). This is consistent with the fact that the stimulation frequency of clinical Vim-DBS is usually chosen to be ≥ 130 Hz [4][5][6]. We observed a short transient with large amplitude in the simulated EMG during Vim-DBS, and the tremor intensity might be higher in the initial ∼ 200 ms after Vim-DBS onset [65][20].

We compared the model simulations with clinical data, which were raw EMG samples recorded from the extensor of an essential tremor patient during DBS-OFF and 135-Hz Vim-DBS (data from Cernera et al. (2021) [10]) (**Figure 5**). The model simulations were presented in steady state (beyond 2 s), and were similar to these clinical data in terms of the EMG amplitudes (**Figure 5**). We observed a 4 – 6 Hz tremor in the clinical EMG sample with DBS-OFF; this is consistent with the corresponding model simulation (**Figure 5**). During 135-Hz Vim-DBS, in both clinical data and model simulation, we observed that the tremor activities are mostly suppressed (**Figure 5**). The tremor is suppressed to a higher extent during 135-Hz Vim-DBS in clinical data compared with the model simulation (**Figure 5**). However, there is high variability in EMG activities across different individuals, and thus the clinical Vim-DBS frequency need to be optimized specifically for individual ET patients [21][10][40].

**Figure 5.**
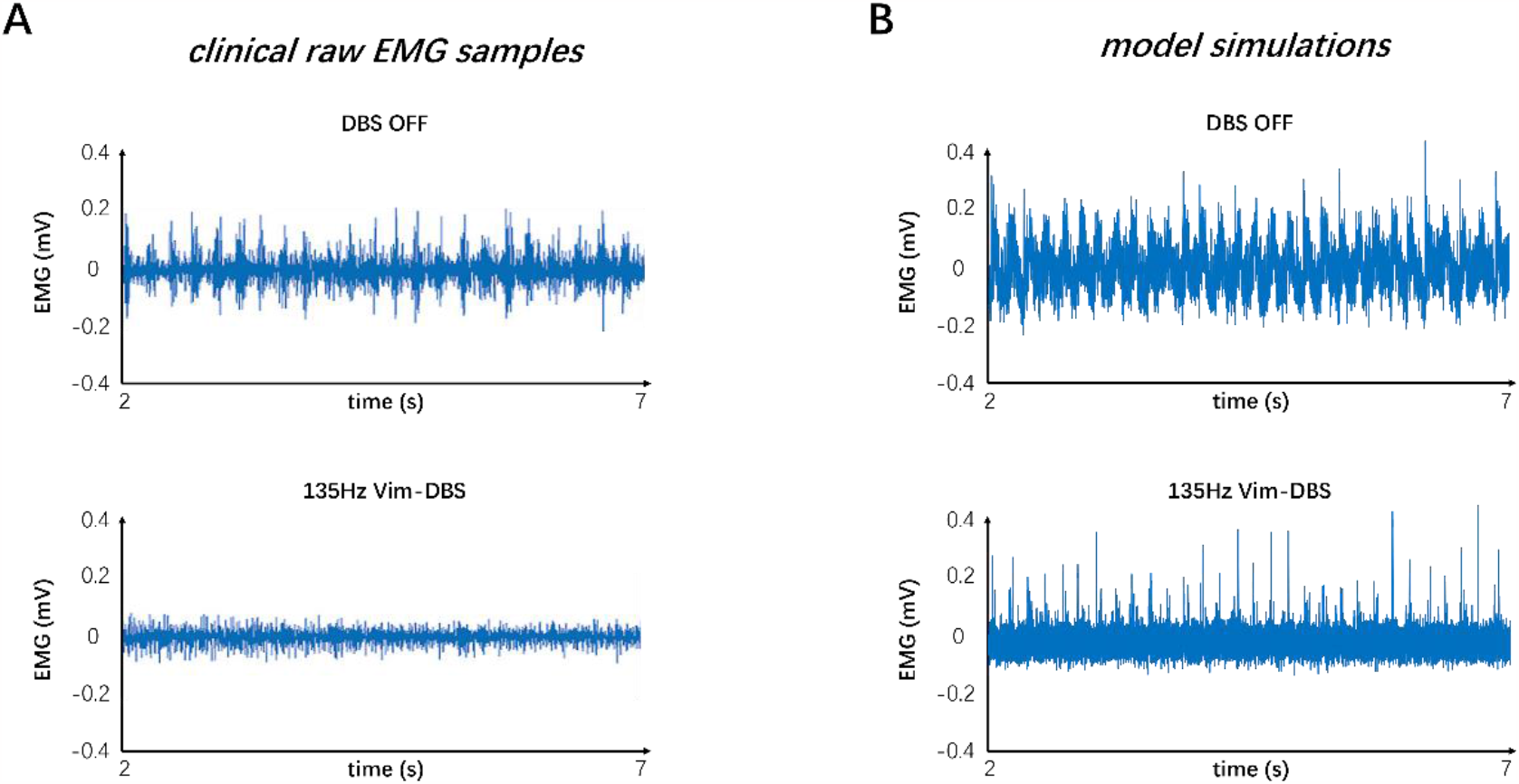
Compare clinical raw EMG samples and model simulations. The clinical raw EMG samples were recorded from the extensor of an essential tremor patient during DBS-OFF and 135-Hz Vim-DBS (data from Cernera et al. (2021) [10]). We compare these clinical data with the steady state EMG simulations from our model.

### Estimation of the Model-Simulated EMG

The EMG simulation from the model is slow for practical implementation. Thus, we estimated the model-simulated EMG with a polynomial fit, to facilitate the computation speed in a closed-loop control system. The model-simulated EMG and polynomial-estimated EMG are denoted as “reference EMG” and “estimated EMG”, respectively (**Figure 6**). We compared the reference EMG and estimated EMG in response to different frequencies of DBS (**Figures 6 and 7**).

**Figure 6.**
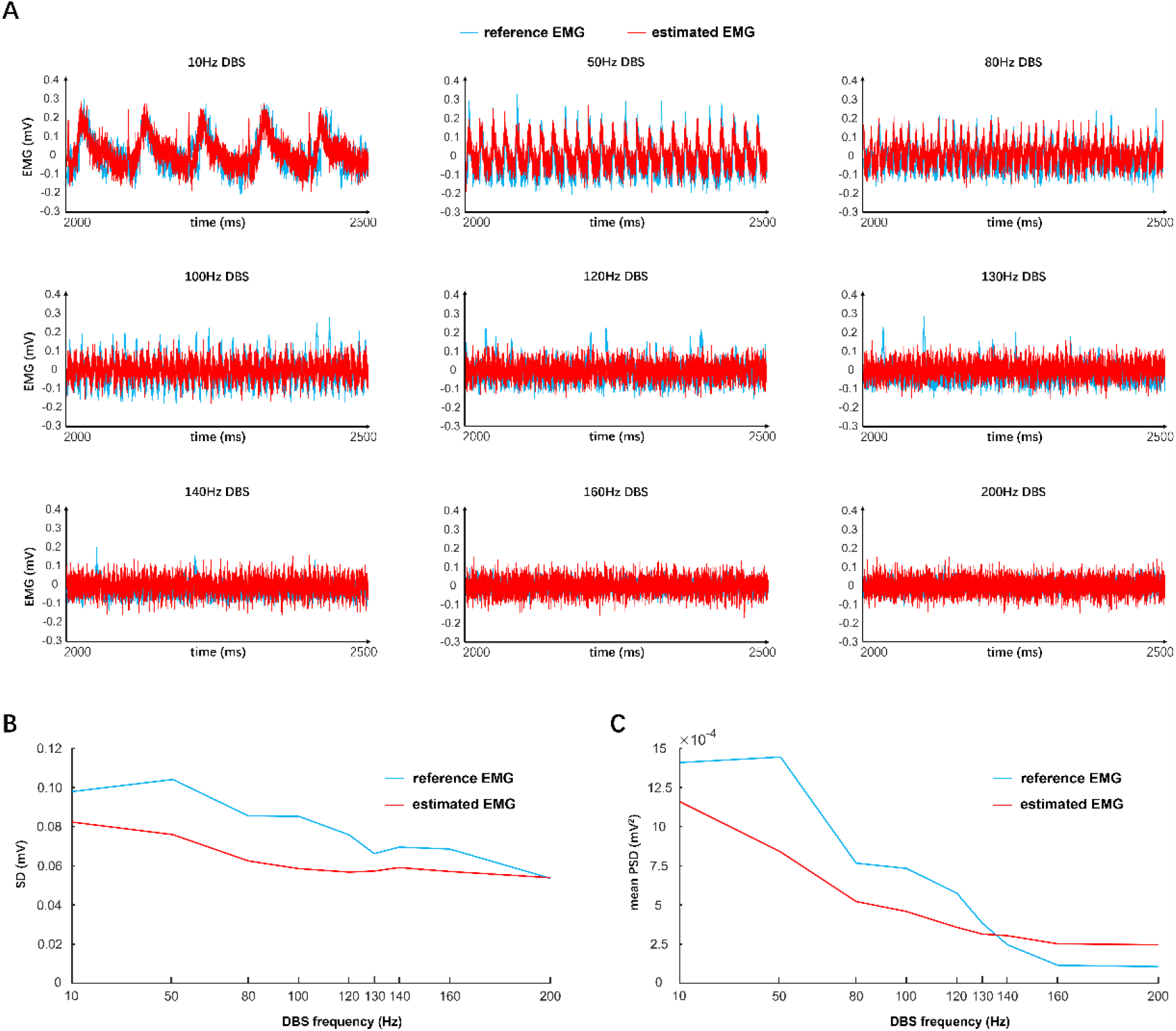
Compare Reference and Estimated EMG (Time Domain) in Response to Different Frequencies of DBS. “Reference EMG” is the EMG simulated by our model (y(t) in **Figure 2**). “Estimated EMG” is the estimation of reference EMG with the polynomial fit (ŷ (t) in **Figure 2**). (A)Compare the reference EMG and estimated EMG in the time domain, in response to different frequencies of DBS. Each signal is subtracted by its mean. (B)Compare the standard deviation (SD) between the reference EMG and estimated EMG. SD is computed based on the initial 5 s of data. (C)Compare the mean power spectral density (PSD) between the reference EMG and estimated EMG. Mean PSD is computed based on the initial 5 s of data.

**Figure 7.**
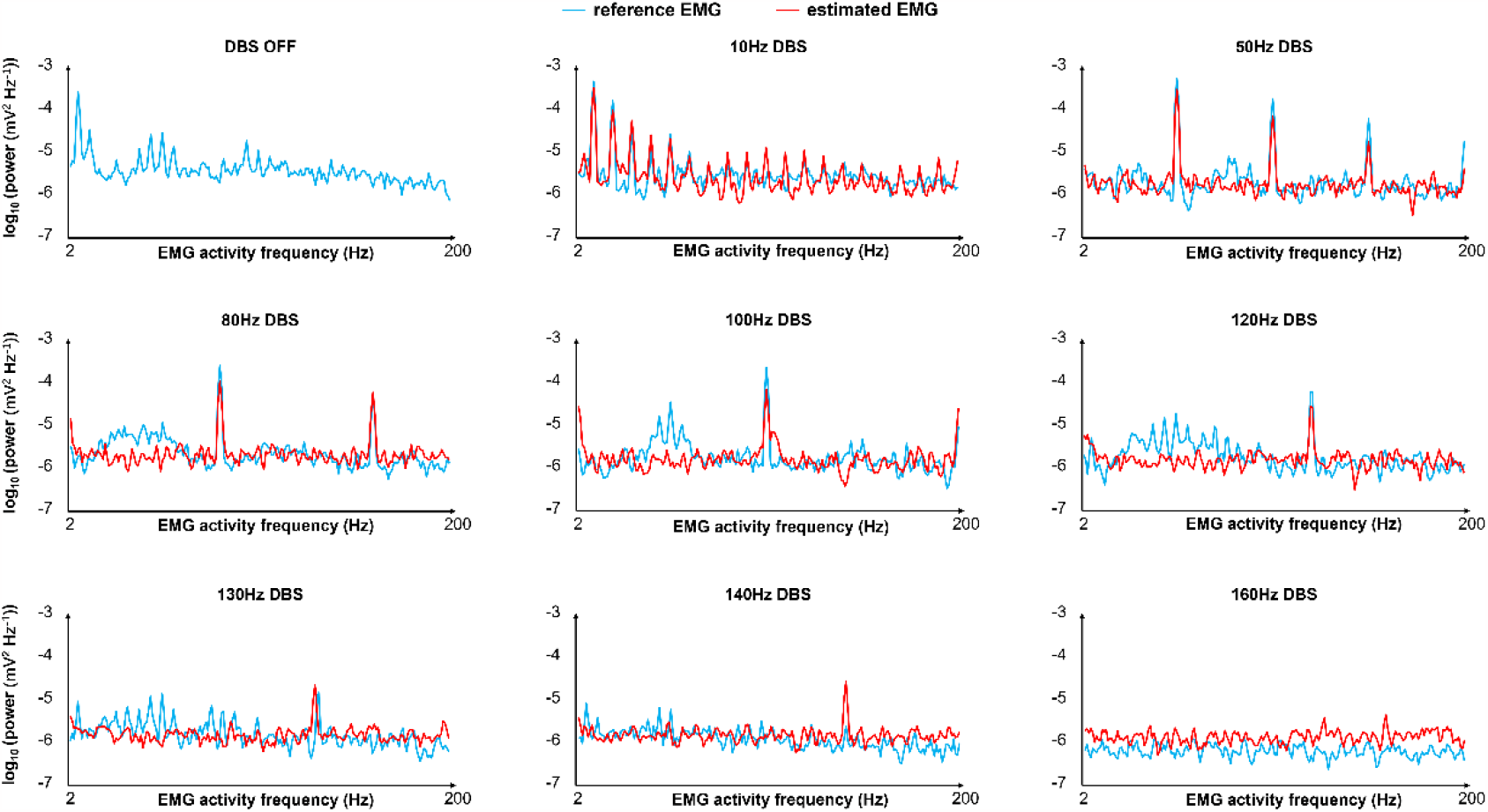
Compare Reference and Estimated EMG (Frequency Domain) in Response to Different Frequencies of DBS. “Reference EMG” is the EMG simulated by our model (y(t) in **Figure 2**). “Estimated EMG” is the estimation of reference EMG with the polynomial fit (ŷ (t) in **Figure 2**). We compare reference EMG and estimated EMG in the frequency domain, which is the frequency band [2, 200] Hz of the EMG activities. For each DBS frequency, at each frequency of the EMG activities, we compute the corresponding frequency power with **Equation 7** in the initial T = 5 s of the EMG. The frequency power of EMG activities is plotted on a log scale.

In the time domain, the estimated EMG is similar to the reference EMG across different DBS frequencies (10 – 200Hz), in terms of both amplitude and variation (**Figure 6**). In particular, in terms of the standard deviation of the initial 5 s of data, the estimated EMG fit the reference EMG well in response to clinical Vim-DBS frequencies (≥ 130 Hz) (**Figure 6B**). Besides the comparison in time domain, we also compared the reference and estimated EMG in frequency domain (**Figure 7**). At each frequency in the band [2, 200] Hz of the EMG activities, we computed the corresponding frequency power with **Equation 7** in the initial T = 5 s of the EMG (**Figure 7**). The estimated EMG (by a 25-order polynomial (**Equation 5**)) is well fitted to the reference EMG in frequency domain, with R^2^ = 0.745 (**Supplementary Figure 3**), computed based on the signals across different DBS frequencies. Additionally, we observed other similarities between the estimated and reference EMG, in terms of the amplitude and pattern of different frequency power of EMG activities (**Figure 7**). The EMG power is high with DBS-OFF and during low frequency (< 100 Hz) DBS, and is mostly suppressed during ≥ 130 Hz DBS (**Figure 7**). During 10 – 80 Hz DBS, in both estimated and reference EMG, we observed that the power is high at the harmonics of the DBS frequency (**Figure 7**). This might indicate that DBS could induce synchronized activities during low frequency of DBS [66][62]. The similarities between the reference and estimated EMG – in both time domain and frequency domain – indicate that the estimated EMG is a proper substitute of the reference EMG for controlling the frequencies of Vim-DBS for treating essential tremor.

### The System Output Based on EMG Power Spectral Density

We computed the system output to be implemented in a closed-loop controller updating the DBS frequency. The system output during Vim-DBS was computed with the estimated EMG (**Equation 5**) that facilitates the implementation speed. For DBS frequency *u*, we defined the system output *z*(*u*) as the mean power spectral density (PSD) in the initial interval *T* = 5 s of the estimated EMG in response to DBS with stimulation frequency *u* (**Equation 9**). PSD represents the full band [2, 200] Hz of EMG activities (**Equation 8**). The system output in response to different frequencies ([10, 200] Hz) of DBS was presented in **Figure 8**:

**Figure 8.**
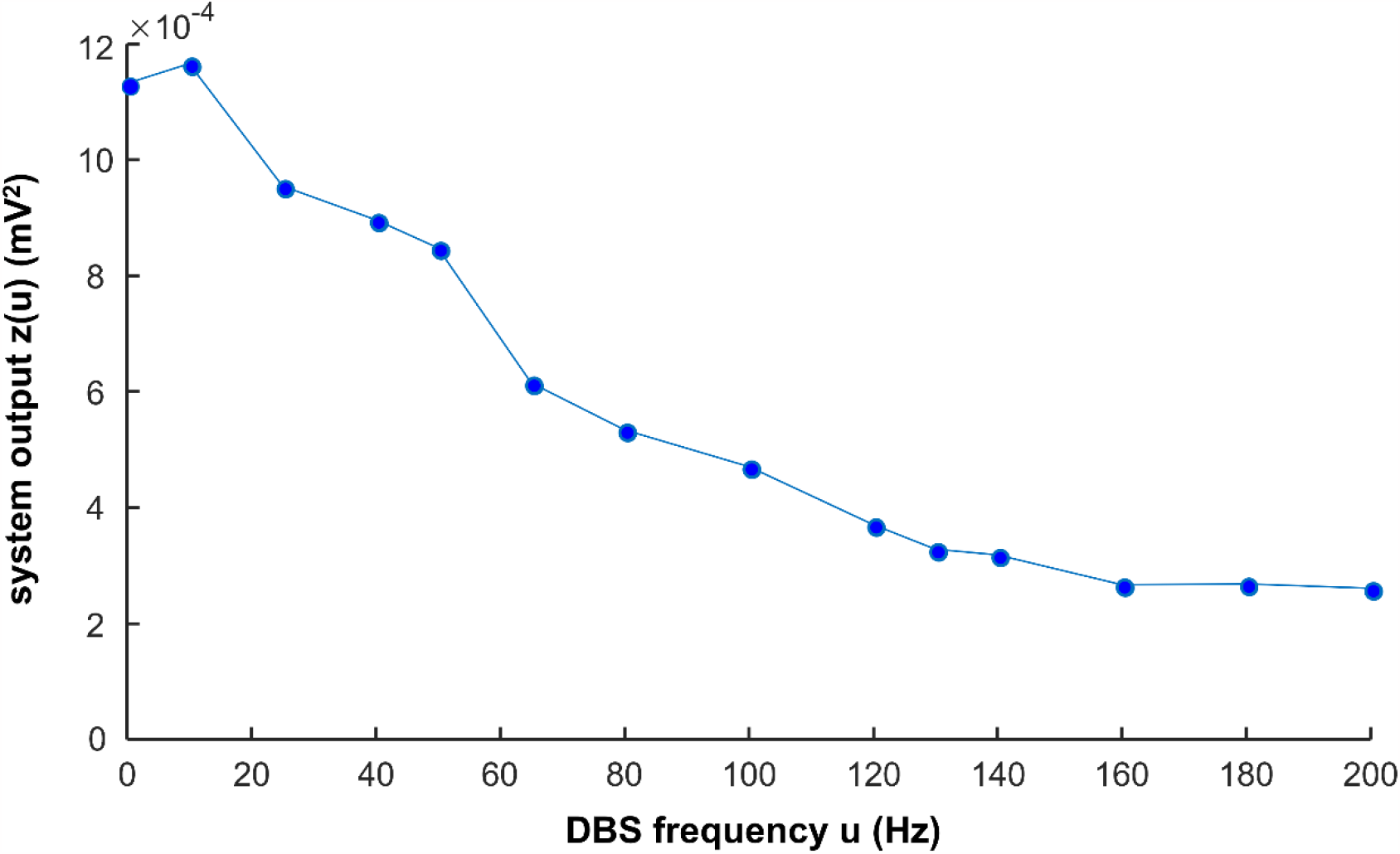
System Output in Response to Different Frequencies of DBS. The system output z(u) is the mean power spectral density (PSD) in the initial interval T = 5 s of the estimated EMG (ŷ (t) in **Figure 2**) in response to DBS with stimulation frequency u (**Equation 9**).

In general, the system output decreases as DBS frequency increases (**Figure 8, Supplementary Table 3**). During low frequency (≤ 50 Hz) DBS, the system output is not much reduced compared with DBS-OFF (*u* = 0) situation (**Figure 8**). The system output is clearly reduced during high frequency (≥ 100 Hz) DBS, and is close to minimum during ≥ 130 Hz DBS (**Figure 8**). These responses of the system output to different DBS frequencies are consistent with clinical observations of the effectiveness of different frequencies of Vim-DBS [62][64][40][4].

### Closed-Loop Control of the DBS Frequency with PID Controller

A PID controller (**Equation 10**) was implemented to update the DBS frequency in closed-loop, based on the system output z (**Equation 9**). The parameters (*K*_*p*_, *K*_*i*_, *K*_*d*_) of the PID controller were chosen to be (10^3^, 10^5^, 5 × 10^3^); parameter tuning was performed to increase the efficiency and robustness of the control (**Supplementary Figure 3** and **Supplementary Note**).

As a test of the controller, when the target DBS frequency is 130 Hz, the PID controller can accurately find the correct DBS frequency in 10 minutes (**Figure 9A** and **Supplementary Table 4**). This shows that our control system is potentially effective and efficient for clinical implementations. Note that during the PID control, only the steady-state DBS frequency (reached after ∼ 10 minutes) is delivered to the patient. As we change the target value of the system output, the result of the PID control is also robustly and flexibly changed (**Figure 9B**). In **Figure 9B**, the five target values of the system output correspond to both trained and untrained (unobserved) DBS frequencies (**Supplementary Table 4**). The trained DBS frequencies ({10, 50, 80, 100, 120, 130, 140, 160, and 200 Hz}) were used in fitting the polynomial coefficients (**Equation 5**), and the untrained (unobserved) DBS frequencies are arbitrary. Our control system is flexible to different targets, and this implies that the control system is potentially flexible to different patients and diseases.

**Figure 9.**
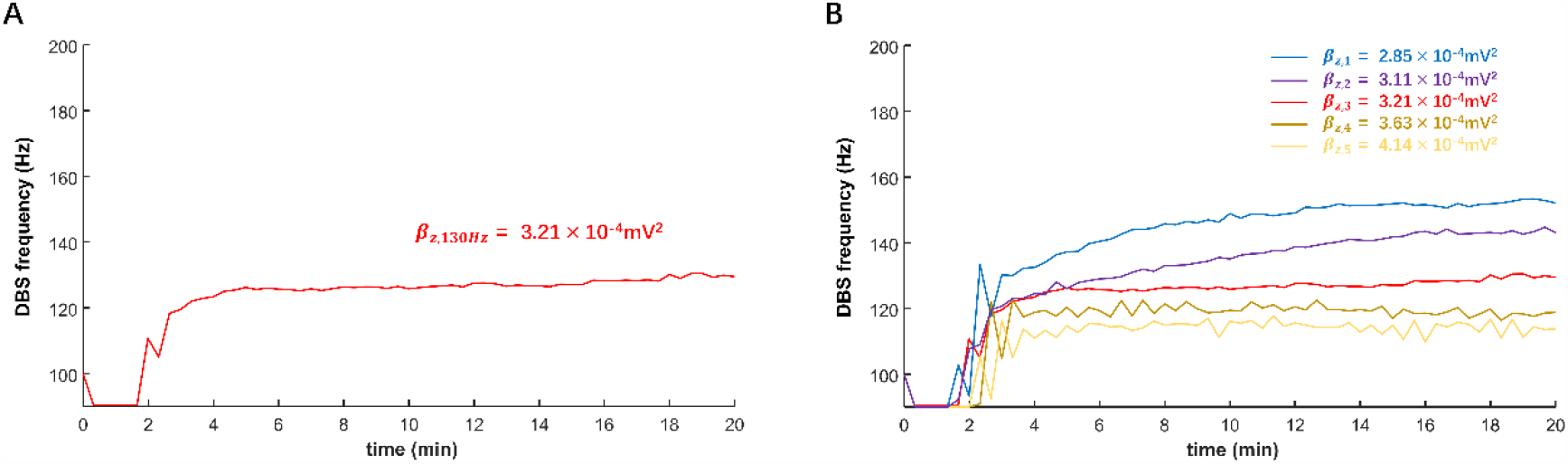
Closed-Loop Control of the DBS Frequency with PID Controller. The DBS frequency is controlled in closed-loop with the proportional-integral-derivative (PID) controller (**Equation 10** and **Figure 2**). The simulation of the PID controller is performed with MATLAB R2022b. (A)The PID control of DBS frequency with target as 130Hz. β_z,130Hz_ is the value of the system output z (**Equation 9**) corresponding to a clinical-DBS frequency = 130 Hz. (B)The PID control of DBS frequency with different targets. β_z,1_, β_z,2_, …, β_z,5_ represent 5 target values of the system output z (**Equation 9** and **Supplementary Table 4**). β_z,2_, β_z,3_ and β_z,4_ correspond to the target DBS frequency 140 Hz, 130 Hz and 120 Hz, respectively. β_z,1_ and β_z,5_ correspond to unobserved DBS frequencies.

## Discussion

We developed a model-based closed-loop control system of the stimulation frequency of Vim-DBS. The DBS control system was based on our previously verified computational model, which represents the neuronal network characterizing the physiological mechanisms that connect the input (DBS pulses) and the output (model-predicted EMG activities). In order to facilitate the implementation speed, we estimated the model-predicted EMG with a polynomial fit, which was used as the feedback biomarker for the controller. The power spectrum of the biomarker was the system output implemented in a PID controller that automatically updates the appropriate DBS frequency. Thus, the closed-loop system controls the EMG power by adjusting DBS frequency. Our closed-loop system can flexibly control the DBS frequency corresponding to different target values of the EMG power, which implies that the control system can potentially be implementable for different diseases and individual patients.

### Clinical Relevance of the System Output

The system output used in our closed-loop system is related to the power of the model-predicted EMG signals and the optimal DBS frequency is obtained by bringing the system output to a specified target value. In clinical studies, the power of EMG is a commonly observed indicator for different movement disorders, e.g., PD [41], ET [38] and akinesia [67]. Tremor symptoms, characterized by the tremor amplitude and frequency, can be identified using the power of EMG [39]. Tremor amplitude is the primary indicator of the severity of tremor [39]. Tremor frequency can be used to partially differentiate disease types; for example, the peak tremor frequency observed in EMG of PD patients is often 3 ∼ 6 Hz [41][39], and in EMG of ET patients is often 4 ∼ 8 Hz [38][21]. Thus, using the power of EMG as a biomarker for ET in a closed-loop DBS is clinically relevant [20][21]. During the DBS control, the target value of the EMG power should be appropriate: a high EMG power indicates the insufficiency of tremor suppression, and a low EMG power can be related to akinesia [67] and myasthenia gravis [68].

### Importance of Predictability in a Control System

The ability to predict how different patterns (e.g., frequency) of DBS change neural- and behavioral-activities is the main advantage of the model-based closed-loop DBS. A controller (e.g., PID) selects appropriate DBS patterns which are biophysically relevant and clinically effective. Although the model parameters (for both encoding- and decoding-models (see **Figure 2**)) are obtained based on sparse DBS frequencies, {5, 10, 20, 30, 50, 100, 130, 200} Hz (human Vim data [33] and non-human primate cortical data [57]), the biophysically-realistic encoding model can robustly predict the effect of an arbitrary DBS frequency in the continuous spectrum of {5 – 200} Hz DBS (see **Figure 4** for some examples; see Table 1 in Tian et al. (2023) [31] for a test of robustness of the firing rate model). Model-based controlling of DBS was addressed in previous computational studies [29][9][28]. Despite the usability of these control systems for in silico explorations, underlying models were not fitted to experimental data. More importantly, these models do not consider physiological mechanisms of the DBS effects. In this work, our (encoding) model not only incorporates biophysically-realistic dynamics of DBS-induced short-term synaptic dynamics but also provides an accurate fit to Vim-DBS experimental data [33]. Additionally, our data-driven decoding model provides a reasonable fit (see **Figures 6 and 7**) to simulated EMG data in which physiological mechanisms of tremor symptoms were preserved (see **Figure 2** for details of simulation study). Our control system predicts the effect of DBS frequency on the power of EMG and delivers optimal DBS frequency.

### The Use of Machine Learning Methods in Closed-Loop DBS

In recent closed-loop DBS control systems, machine learning methods have been developed to map biomarker features (input) to patients’ observed states (output) and could further deliver an appropriate DBS setting [25][69][70]. Therefore, it is imperative to identify key biomarkers that need to be extracted from the LFP and EMG as they will serve as input features for a judiciously selected machine learning model. Castaño-Candamil et al. (2020) [26] used a regression method to estimate tremor severity from electrocorticographic (ECoG) power in ET patients, and adjusted DBS intensity according to tremor severity. Golshan et al. (2018) [71] utilized the wavelet coefficients of STN-LFP beta frequency range as features and further developed a support vector machine (SVM) classifier for the behaviors of PD patients. Numerous prior studies have established high performance using SVM classifiers with input features such as phase amplitude coupling [25], Hjorth parameters [72], beta band power [25], and burst duration [70]. Using power densities within the beta [69] and gamma [73] bands as features, Hidden Markov Models [70][74][73], SVM [71], convolutional neural networks (CNN) [70][75][72], linear discriminant analysis (LDA) [70], and logistic regression [76] were performed. It was also recommended in a couple studies that deep learning methods such as CNNs are worth investigating as they capture nonlinear temporal dynamics and waveform shape [70][72]. For example, Haddock et al. (2019) [27] developed a deep learning method that classified the behaviors of ET patients based on the PSD of ECoG and used the classification results to turn DBS ON/OFF.

A key improvement to existing machine learning methods is to incorporate physiological characterizations of the input-output mapping. In this work, we developed a physiological model to map the input (DBS frequency) to the output (model-predicted EMG). We then used a polynomial-based approximation to estimate the input-output map to facilitate the implementation speed of the control system. However, a polynomial method is prone to be less robust to unseen inputs, due to its high-order terms [77][78]. Thus, an important line of future works is to use state-of-the-art machine learning methods, particularly deep learning methods, to replace the simple polynomial-based input-output mapping. Consequently, it is important to understand key EMG features for muscle activation in Parkinson’s disease. Literature highlights sample kurtosis, recurrence rate, and correlation dimension as three specific EMG features that are responsive to changes in DBS-treatment parameters [79]. These features have been fed as input into LDA, CNN, and SVM, where SVM performed the best [80]. In a few investigations, EMG features, encompassing frequency, amplitude, and regularity, were scrutinized [81][82]. The signal mean and power of the peak frequency performed well as features when using a random forest model and a deep learning network for adaptive DBS [81]. It is important to note that while simpler models like LDA are valued [83][84] for their interpretability in the context of DBS for PD, the limited availability of labeled datasets resulted in ambiguous success for complex models, particularly deep neural networks [72].

Our closed-loop DBS control is based on a physiological model that can generate an arbitrary amount of synthetic data, which can be implemented in fully training deep learning methods. The arbitrary amount of data in the training set will increase the accuracy and robustness of our future closed-loop DBS control based on physiological models and deep learning methods. Therefore, it is recommended to evaluate the efficacy of SVM, logistic regression, LDA, hidden Markov Model (HMM), random forests, and deep neural network models like CNNs in greater detail using the abundance of synthetic data. Watts et al. (2020) [84] performed a thorough retroactive study of various machine learning classifiers used to identify optimal DBS parameters for PD, and it was recommended to pursue machine learning in the context of adaptive closed-loop DBS for PD.

### Limitations and Future Work

Our model-based closed-loop system included the clinical Vim-DBS data recorded in Vim neurons across different stimulation frequencies (10 – 200 Hz) [33]. However, experimental data were not sufficiently included in other components of the closed-loop system. We incorporated non-human primate single-unit recordings in M1 during 130-Hz VPLo-DBS reported in Agnesi et al. (2015) [57], but M1 activities in response to other DBS frequencies were not recorded. The EMG model simulation is consistent with some clinical observations, but was not further developed and validated by fitting clinical EMG data. In fact, in current experimental works of DBS, EMG signal is usually recorded with DBS-OFF or high DBS frequencies (> 100 Hz) [21][40][85], and there lacks EMG recordings in response to a wide spectrum of DBS frequencies (e.g., 10 – 200 Hz). In the future, we plan to incorporate more experimental data into the further development of the model-based closed loop DBS control, and these experimental data – in particular, cortical and EMG data – need to be recorded with different DBS frequencies from each individual subjects.

Worth mentioning is that the synaptic connections between the Vim (VPLo in primate) are reciprocal and excitatory, though the projections that the M1 sends to the Vim are in different M1 laminae than the ones it receives from Vim [86]. Our model simplified this excitatory feedback relationship by considering the excitatory propagation effect as uni-directional, from the Vim to the M1. We will develop more detailed Vim-M1 model in the future. Our future closed-loop DBS systems will be constructed based on both improved models and deep learning methods.

