## Supplementary Method, Tables, and Figures for "Model-Based Closed-Loop Control of Thalamic Deep Brain Stimulation"

### Supplementary Method – The Tsodyks & Markram Model of Short-Term Synaptic Plasticity

The synapses of the neuron in the primary motor cortex (M1) were characterized by the Tsodyks & Markram (TM) model of short-term synaptic plasticity (STP) [1]. During Vim-DBS, we modeled that each M1 neuron receives DBS-induced inputs from 500 synapses, with 90% excitatory synapses ($N_{exc}$ = 450) and 10% inhibitory synapses ($N_{inh}$ = 50) ([2], **Table S1**). The DBS-induced inputs were from two Sources: (1) The direct DBS activation of the axons projected to the M1 neuron; and (2) The firings of the Vim neurons during Vim-DBS. In Source (1), we assumed that each DBS pulse generates a spike in each of the 500 synapses simultaneously [2][3]. In Source (2), from our previous Vim-network model [ref], we simulated the instantaneous firing rate of the Vim neurons receiving DBS of different stimulation frequencies (10~200Hz). Then, the Vim firing rate signal was implemented as the time-varying Poisson rate to generate Poisson spike trains. The spikes from Sources (1) and (2) were passed to the TM model (**Equations 2-4**) to generate the DBS-induced post-synaptic current $I_{DBS}$.

Besides the DBS-induced inputs, the M1 neuron also received the background neuronal spikes that induce the tremor. We modeled the tremor-inducing background firing rate as a waveform consisting of 6Hz bursts and a baseline shift, each burst consists of 3 consecutive sinusoidal waves, and the period of each wave is 20ms (**Supplementary Figure 1**). We then generated Poisson spike trains from the background firing rate waveform, and these spikes were passed to TM model (**Equations 2-4**) to generate the post-synaptic current $I_b$ induced by these background spikes generating the tremor. $I_{syn} = I_{DBS} + I_b$ is the total post-synaptic input current that incorporates all the input spikes.

$I_{syn}$ was obtained by a linear combination of post-synaptic excitatory ($I_{exc}$) and inhibitory ($I_{inh}$) currents as follows:

$$I_{syn}(t) = w_{exc}\, I_{exc}(t) - w_{inh} I_{inh}(t) \qquad \text{(Equation 1)}$$

where $w_{exc}$ and $w_{inh}$ denote the scaling weights of the excitatory and inhibitory currents, respectively (**Supplementary Table 1**). $I_{exc}$ (respectively, $I_{inh}$) is the total post-synaptic current from all excitatory (respectively, inhibitory) synapses; each synapse (excitatory or inhibitory) was modeled by the TM model for short-term synaptic plasticity:

$$\frac{du}{dt} = -\frac{u}{\tau_{facil}} + U(1-u^-)\delta(t-t_{sp}) \quad \text{(Equation 2)}$$

$$\frac{dm}{dt} = \frac{1-m}{\tau_{rec}} - u^+m^-\delta(t-t_{sp}) \quad \text{(Equation 3)}$$

$$\frac{dI}{dt} = -\frac{I}{\tau_s} + Au^+r^-\delta(t-t_{sp}) \quad \text{(Equation 4)}$$

where $u$ is a utilization parameter, indicating the fraction of neurotransmitters ready for release into the synaptic cleft (due to calcium ion flux in the presynaptic terminal). The variable $m$ indicates the fraction of resources remaining available after the neurotransmitter depletion caused by neuronal spikes. We denote as $u^-$ and $m^-$ the corresponding variables just before the arrival of the spike; similarly, $u^+$ and $m^+$ refer to the moment just after the spike. The $\delta$–function models the abrupt change upon the arrival of each presynaptic spike $t_{sp}$; for example, at $t = t_{sp}$ in **Equation 2**, $u$ increases by $U(1-u^-)$, and $\delta(t-t_{sp}) = 0$ when $t \neq t_{sp}$. If there is no presynaptic activity (spike), $u$ exponentially decays to zero; this decay rate is the facilitation time constant, $\tau_{facil}$ (**Equation 2**). In contrast to the increase of $u$ upon the arrival of each presynaptic spike, $m$ drops and then recovers to its steady state value ($=$ 1); this recovery rate is given by the recovery time constant $\tau_{rec}$ (**Equation 3**). The competition between the facilitation ($\tau_{facil}$) and recovery ($\tau_{rec}$) time constants determined the dynamics of the synapse. In the TM model, $U$, $\tau_{facil}$, and $\tau_{rec}$ were the parameters that determine the three types of the synapse: facilitation ("F"), pseudo-linear ("P"), and depression ("S") ([2], **Supplementary Table 1**). In **Equation 4**, $I$ is the post-synaptic current, $A$ is the absolute response amplitude, and $\tau_s$ is the post-synaptic time constant (**Supplementary Table 1**). We obtained $I_{exc}$ (respectively, $I_{inh}$) by adding the post-synaptic currents from all excitatory (respectively, inhibitory) synapses. The TM model parameters in **Supplementary Table 1** were chosen based on the previous modeling works on specific experimental datasets [2][4][5].

### Supplementary Note – Parameter Tuning of the PID Controller

We manually tune the parameters ($K_p$, $K_i$, $K_d$) of the PID controller. The purpose of the PID parameter tuning was to increase the efficiency and robustness of the control. The decisions are ($K_p$, $K_i$, $K_d$) = ($10^3$, $10^5$, $5\times10^3$). For each of the three PID parameter, we compared our decision with other values in a control experiment (i.e., the other two parameters are the same as our decisions)

(**Supplementary Figure 2**). Our PID parameter decisions are compared with other values in the situation where the target DBS frequency is 130Hz (**Supplementary Figure 2A**). The results from the PID controller were constrained to be in the DBS frequency range [10, 200]Hz, and this wide frequency range contributes to the full investigation of the behaviors of the PID controllers.

In the tuning of $K_p$ (**Supplementary Figure 2B**), we observed that the results are non-robust when $K_p \leq 10^2$ or $K_p \geq 5\times10^4$. The results with $K_p = 10^4$ is similar to our decision ($K_p = 10^3$), but the predicted DBS frequency is less accurate (**Supplementary Table 5**). In the tuning of $K_i$ (**Supplementary Figure 2C**), we observed that the results are mostly robust for the shown values of $K_i$. However, the results with the $K_i$ values other than our decision ($K_i = 10^5$) are less accurate ($K_i \geq 2\times10^5$) or less efficient ($K_i \leq 5\times10^4$). In the tuning of $K_d$ (**Supplementary Figure 2D**), we observed that the result with our decision ($K_d = 5\times10^3$) is more robust than the results with other $K_d$ values.

### Supplementary Tables

| *excitatory synapses* $N_{exc}$ = 450; $w_{exc}$ = 49.00 | | | | | | *inhibitory synapses* $N_{inh}$ = 50; $w_{inh}$ = 73.08 | | | | | |
|---|---|---|---|---|---|---|---|---|---|---|---|
| **para** / **type** | $U$ | $\tau_{facil}$ (ms) | $\tau_{rec}$ (ms) | $\tau_s$ (ms) | $A$ | **para** / **type** | $U$ | $\tau_{facil}$ (ms) | $\tau_{rec}$ (ms) | $\tau_s$ (ms) | $A$ |
| **F (40%)** | 0.19 | 670 | 138 | | | **F (40%)** | 0.016 | 376 | 45 | | |
| **P (20%)** | 0.45 | 326 | 329 | 2 | 1 | **P (30%)** | 0.29 | 62 | 144 | 8.5 | 1 |
| **S (40%)** | 0.04 | 17 | 85 | | | **S (30%)** | 0.25 | 21 | 706 | | |

**Supplementary Table 1. Tsodyks & Markram Model Parameters for Synapses Projected to one M1 Neuron**

*We show the parameters related to the Tsodyks & Markram model* [1], *which is implemented to compute the post-synaptic current ($I_{syn}$ , **Equation 1**) into the neuron in the primary motor cortex (M1). For one M1 neuron, we model that it receives inputs from 500 synapses, with 90% excitatory synapses ($N_{exc}$ = 450) and 10% inhibitory synapses ($N_{inh}$ = 50). $w_{exc}$ and $w_{inh}$ are the scaling weights of the post-synaptic excitatory ($I_{exc}$) and inhibitory ($I_{inh}$) currents (**Equation 1**). Both excitatory and inhibitory synapses consist of 3 types: facilitation ("F"), pseudo-linear ("P"), and depression ("S"). For excitatory synapses, "F (40%)" represents "the facilitation type of synapses account for 40% of all the excitatory synapses"; similar meanings for other synaptic types, and the inhibitory synapses. "para" represents "the Tsodyks & Markram model parameters", which consist of "$U$" (scaling factor), "$\tau_{facil}$" (facilitation time constant), "$\tau_{rec}$" (recovery time constant), "$\tau_s$" (post-synaptic time constant) and "$A$" (absolute response amplitude) (**Equations 2 – 4**).*

| | |
|---|---|
| $\varphi_0$ | 0.119 |
| $\varphi_1$ | 0.0891 |
| $\varphi_2$ | 2.12 |
| $\varphi_3$ | 4.26 |
| $\varphi_4$ | -21.3 |
| $\varphi_5$ | -29.0 |
| $\varphi_6$ | 97.8 |
| $\varphi_7$ | 61.0 |
| $\varphi_8$ | -256 |
| $\varphi_9$ | 7.15 |
| $\varphi_{10}$ | 369 |
| $\varphi_{11}$ | -208 |
| $\varphi_{12}$ | -230 |
| $\varphi_{13}$ | 295 |
| $\varphi_{14}$ | -31.3 |
| $\varphi_{15}$ | -136 |
| $\varphi_{16}$ | 97.7 |
| $\varphi_{17}$ | -12.7 |
| $\varphi_{18}$ | -20.7 |
| $\varphi_{19}$ | 16.2 |
| $\varphi_{20}$ | -6.37 |
| $\varphi_{21}$ | 1.59 |
| $\varphi_{22}$ | -0.264 |
| $\varphi_{23}$ | 0.0284 |
| $\varphi_{24}$ | $-1.80\times10^{-3}$ |
| $\varphi_{25}$ | $5.16\times10^{-5}$ |

**Supplementary Table 2. Coefficients of the Polynomial Fit (Equation 5) (3 significant digits)**

| DBS frequency u (Hz) | system output z(u) ($10^{-4}mV^2$) |
|---|---|
| 0 | 11.31 |
| 10 | 11.66 |
| 25 | 9.53 |
| 40 | 8.94 |
| 50 | 8.45 |
| 65 | 6.10 |
| 80 | 5.29 |
| 100 | 4.65 |
| 120 | 3.63 |
| 130 | 3.21 |
| 140 | 3.11 |
| 160 | 2.59 |
| 180 | 2.60 |
| 200 | 2.53 |

**Supplementary Table 3. System output in Response to Different Frequencies of DBS**

*The system output z(u) is the mean power spectral density (PSD) in the initial T = 5s of the estimated EMG ($\hat{y}(t)$ in **Figure 2**) in response to DBS with stimulation frequency = u (Hz) (**Equation 9**).*

| target \ time | 5min | 10min | 15min | 20min |
|---|---|---|---|---|
| $\boldsymbol{\beta_{z,1}}$ | 137.2Hz | 148.8Hz | 151.8Hz | 152.0Hz |
| $\boldsymbol{\beta_{z,2}}$ = z(140) | 126.0Hz | 135.0Hz | 141.7Hz | 143.0Hz |
| $\boldsymbol{\beta_{z,3}}$ = z(130) | 125.9Hz | 125.5Hz | 126.7Hz | 129.2Hz |
| $\boldsymbol{\beta_{z,4}}$ =z(120) | 120.4Hz | 119.4Hz | 117.1Hz | 118.8Hz |
| $\boldsymbol{\beta_{z,5}}$ | 114.7Hz | 116.1Hz | 114.9Hz | 113.8Hz |

**Supplementary Table 4. Updated DBS Frequency during PID Control with Different Targets**

*"PID" represents the proportional-integral-derivative controller (**Equation 10**).* $\beta_{z,1}, \beta_{z,2}, \dots, \beta_{z,5}$ *represent 5 target values of the system output z (**Figure 9B** and **Equation 9**). The values of* $\beta_{z,1}, \beta_{z,2}, \dots, \beta_{z,5}$ *were specified in the legend of **Figure 9B**. This table shows the updated DBS frequency in the process of the PID control, with different target values of the system output.* $\beta_{z,2}$, $\beta_{z,3}$, *and* $\beta_{z,4}$ *are system output target values with target DBS frequency 140Hz, 130Hz and 120Hz, respectively (**Supplementary Table 3**).* $\beta_{z,1}$ *and* $\beta_{z,5}$ *are system output target values corresponding to unknown DBS frequencies, to be explored by the PID controller.*

| **time** / $K_p$ | **5min** | **10min** | **15min** | **20min** |
|---|---|---|---|---|
| **$10^2$** | 200.0Hz | 187.2Hz | 167.9Hz | 149.4Hz |
| **$10^3$ (selected)** | 122.7Hz | 127.7Hz | 129.4Hz | 130.2Hz |
| **$10^4$** | 121.0Hz | 125.3Hz | 126.6Hz | 128.0Hz |
| **$5\times10^4$** | 127.3Hz | 126.1Hz | 129.7Hz | 126.7Hz |
| **$10^5$** | 124.2Hz | 121.6Hz | 123.6Hz | 127.2Hz |

**Supplementary Table 5. Updated DBS Frequency during PID Control with Different $K_p$ Values**

*"PID" represents the proportional-integral-derivative controller, and* $K_p$ *denotes the proportional gain of the PID controller (**Equation 10**). This table shows the updated DBS frequency in the process of the PID control with target DBS frequency 130Hz, using different values of* $K_p$. *The corresponding PID control processes are presented in **Supplementary Figure 2B**. The selected* $K_p$ *value is* $10^3$ *(**Supplementary Figure 2A**).*

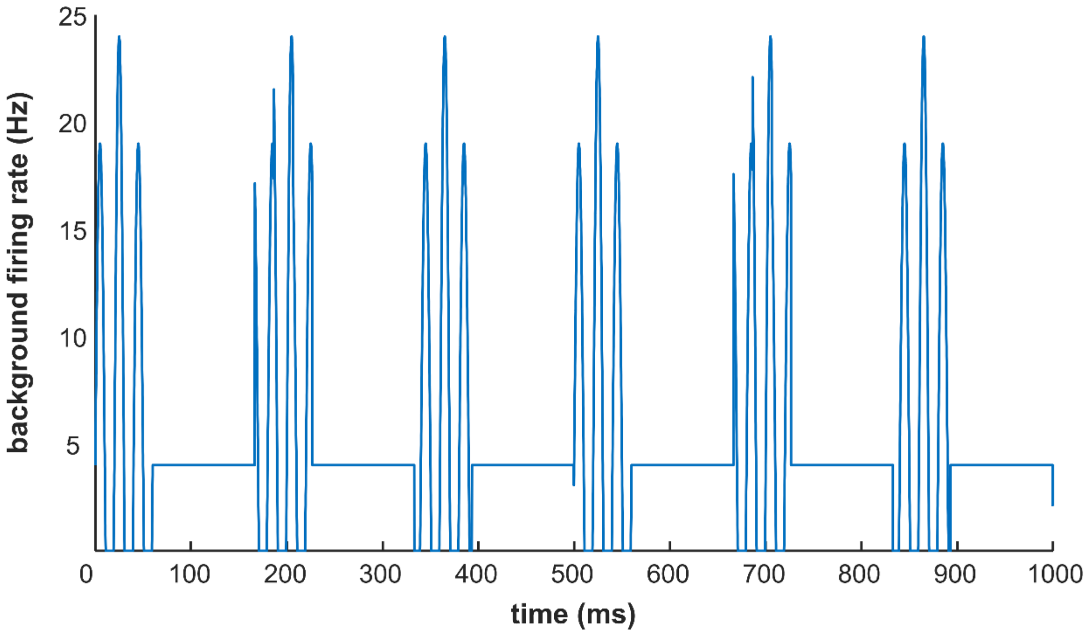


**Supplementary Figure 1. Background Cortical Inputs that Induce Essential Tremor**

*The background inputs into the neuron in the primary motor cortex (M1) induce essential tremor symptoms, which are often in the frequency band 4~8Hz. We model these background inputs as a firing rate waveform consisting of 6Hz bursts and a baseline shift. Each burst consists of 3 consecutive sinusoidal waves and the period of each wave is 20ms.*

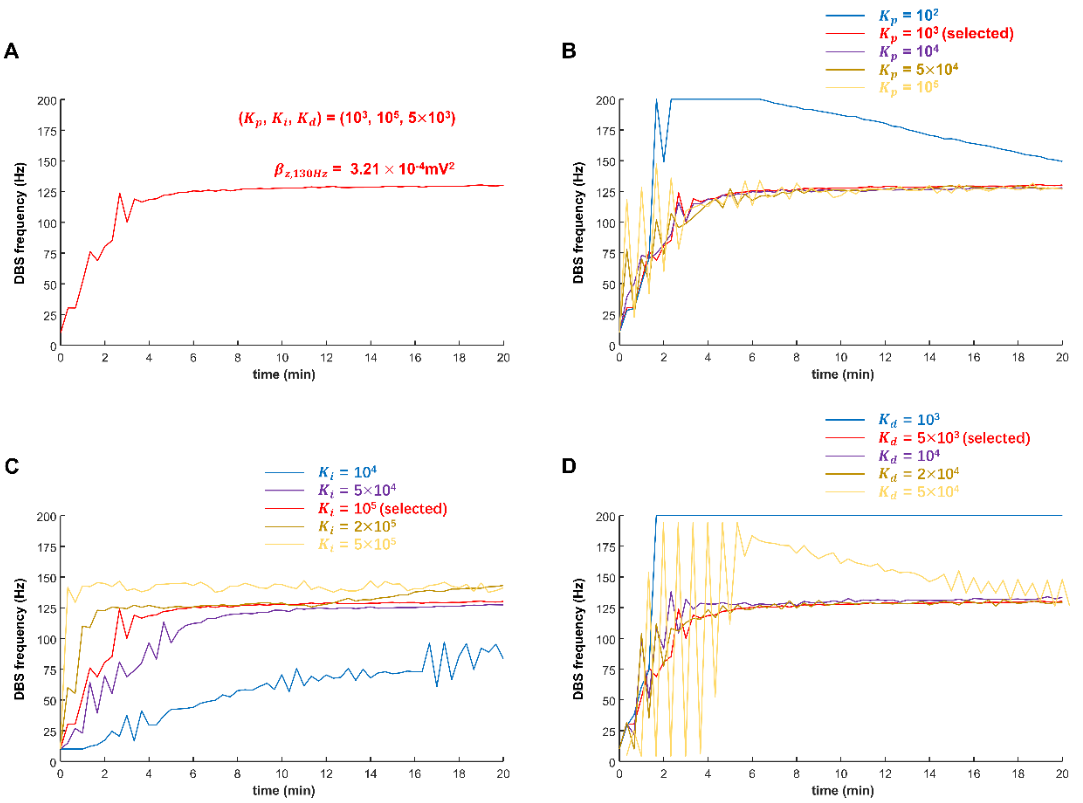


**Supplementary Figure 2. Parameter Tuning of the PID Controller**

*We compare the impact of different parameters ($K_p$, $K_i$, $K_d$) of the proportional-integral-derivative (PID) controller (**Equation 10**). We test the parameters ($K_p$, $K_i$, $K_d$) of PID control with the target DBS frequency 130Hz. The results from the PID control are constrained to be in the DBS frequency range [10, 200]Hz.*

*(A) Our decisions of the parameters ($K_p$, $K_i$, $K_d$), and the PID control process corresponding to these selected parameters. $\beta_{z,130Hz}$ is the value of the system output z (**Equation 9**) corresponding to DBS frequency = 130Hz.*

*(B) The PID control processes corresponding to different $K_p$. The other two parameters ($K_i$, $K_d$) are the same as our decisions in (A), in all the PID control processes.*

*(C) The PID control processes corresponding to different $K_i$. The other two parameters ($K_p$, $K_d$) are the same as our decisions in (A), in all the PID control processes.*

*(C) The PID control processes corresponding to different $K_d$. The other two parameters ($K_p$, $K_i$) are the same as our decisions in (A), in all the PID control processes.*

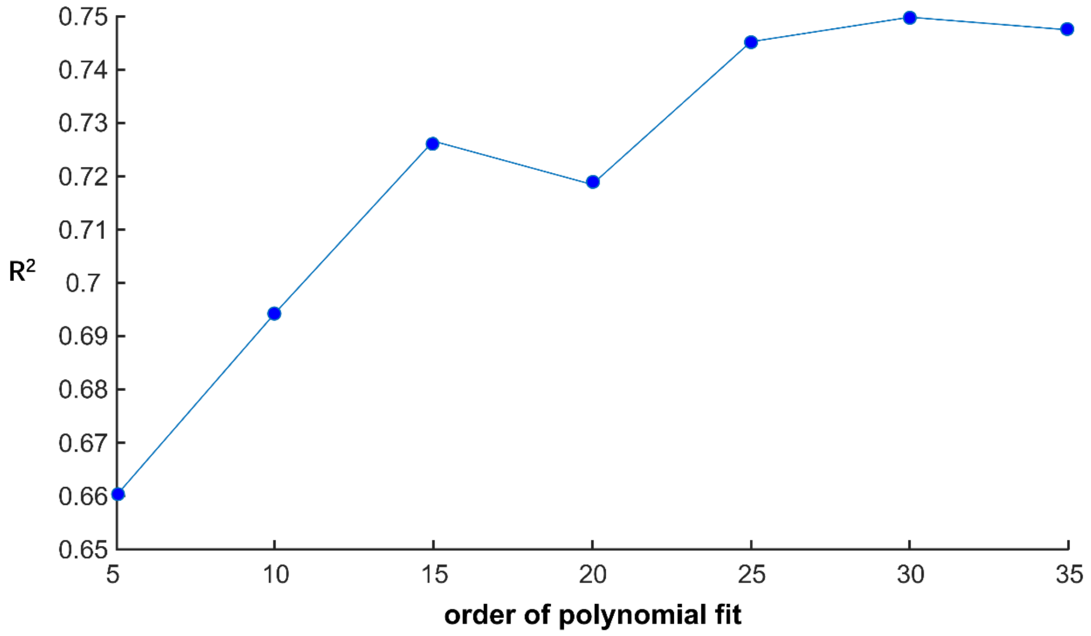


**Supplementary Figure 3. Comparison of $R^2$ of Different Orders of Polynomial Fits Using Frequency Domain Signals**

We compare the $R^2$ statistic of different orders of polynomial fits using the corresponding frequency domain signals across different DBS frequencies (illustrated in **Figure 7**). The data length is 10s for each DBS frequency in {10, 50, 80, 100, 120, 130, 140, 160, and 200Hz}.
